# Genetic evidence challenges the native status of a threatened freshwater fish (*Carassius carassius*) in England

**DOI:** 10.1101/026088

**Authors:** Daniel L Jeffries, Gordon H Copp, Lori Lawson Handley, Carl D Sayer, Bernd Hänfling

## Abstract

A fundamental consideration for the conservation of a species is the extent of its native range, however defining a native range is often challenging as changing environments drive shifts in species distributions over time. The crucian carp, *Carassius carassius* (L.) is a threatened freshwater fish native to much of Europe, however the extent of this range is ambiguous. One particularly contentious region is England, in which *C. carassius* is currently considered native on the basis of anecdotal evidence. Here, we use 13 microsatellite loci, population structure analyses and approximate bayesian computation (ABC), to empirically test the native status of *C. carassius* in England. Contrary to the current consensus, ABC yields strong support for introduced origins of *C. carassius* in England, with posterior distribution estimates placing their introduction in the 15th century, well after the loss of the doggerland landbridge. This result brings to light an interesting and timely debate surrounding our motivations for the conservation of species. We discuss this topic, and make arguments for the continued conservation of *C. carassius* in England, despite its non-native origins.

## Introduction

Obtaining a detailed understanding of a species’ native range and the distribution of its diversity within that range is fundamental for species conservation (Frankham *et al.* 2002; Reed & Frankham 2003; Scoble & Lowe 2010; IUCN 2012). However, this is complicated by the fact that species’ ranges are not static but often change dramatically over time in response to changing environments and newly arising dispersal corridors. A species is usually considered native if it has colonised an area naturally whereas areas which have been colonised without human intervention are not included as part of the native range which has profound implications for management (e.g. Copp et all 2005). During the last 2.5 MY, the ranges of European biota have been impacted most strongly by the glacial cycles (Hewitt 1999). These processes have been extensively studied in particularly in freshwater fish, whose postglacial recolonisation dynamics have been determined by the history of river drainage systems (Bianco 1990; Bănărescu 1990, 1992; Bernatchez & Wilson 1998; Reyjol *et* al. 2006). For example, ephemeral rivers and periglacial lakes that result from glacial meltwater have provided opportunities for fish colonisations (Gibbard *et al.* 1988) of otherwise isolated drainages (Grosswald 1980; Arkhipov *et al.* 1995). However, human-mediated translocations also had a significant impact on the current distributions of European freshwater fish have also been determined, which have enabled some species to overcome natural dispersal barriers like watersheds (Copp *et al.* 2005; Gozlan *et al.* 2010). Knowing whether natural or human mediated dispersal, is responsible for an organism’s contemporary distribution, is fundamental in determining its native range.

However, this distinction is particularly difficult to make in the UK. With very few exceptions such as groundwater invertebrates (McInerney *et al.* 2014), it is thought that the vast majority of terrestrial and freshwater animals were forced South, into continental refugia, by the expansion of the Weichselian ice sheet during the last glaciation. At its maximum extent, approximately 25000 years before present (YBP), this ice sheet covered almost the entirety of the UK, with frozen tundra covering the remaining unglaciated land area (Coles 2000). Native UK species have therefore recolonised this region over the last 18,000 years, when the Weichselian ice sheet began to recede. In the case of primary freshwater fish, this was made possible by connections between English and Continental river systems that existed in Doggerland, the land bridge connection between southeast England and continental Europe. However, this window of opportunity was relatively short, as Doggerland was inundated at around 7800 YBP with rising sea levels resulting from the continued melting of the Weichselian ice sheet (Coles 2000).

After the loss of the Doggerland land bridge, the only means by which freshwater species could colonise the UK, precluding the very unlikely possibility of fertilised eggs being transported by migrating waterfowl (for which no empirical evidence exists, to our knowledge), would have been via human mediated introductions. The earliest known record of live fish translocations into the UK was the movement of common carp, *Cyprinus carpio*, into the southeast of England by monks in the 15^th^ century (Lever 1977). Although, it cannot be ruled out that they were introduced by earlier civilisations, e.g. the Romans, in the 1^st^ century A.D or in the following few centuries by Viking invaders.

The dates described above therefore allow us to make a clear distinction between the possible arrival times of a primary freshwater fish in the UK under two hypotheses; if native, then it must have colonised naturally before 7800 YBP, if introduced, then realistically it could not have arrived earlier than approximately 2000 YBP.

One species, which, in the past, has had a particularly contentious status in the UK is the crucian carp (*Carassius carassius*, Linneaus 1758); a primary freshwater fish, native to much of central and Eastern Europe. The crucian carp is of conservation concern in much of its range due to sharp declines in the number and sizes of populations in recent times, which has led to local population extinctions (Copp *et al.* 2010; Savini *et al.* 2010; Sayer *et al.* 2011; Mezhzherin *et al.* 2012; Rylková *et al.* 2013). Awareness of the threats to *C. carassius* is building, and it often appears on red-lists at the national level e.g, Czech Republic (Lusk *et* al. 2004), Ukraine (Andrievskiy 2009), Austria (Wolfram & Mikschi 2007), Croatia (Mrakovčić *et al.* 2007) and Serbia (Simic, V *et al.* 2009). Despite this, however, there are still very few active conservation initiatives for *C. carassius* in Europe and, to our knowledge, one of the most comprehensive of these exists in Norfolk, in eastern England (Copp & Sayer 2010; Sayer *et al.* 2011).

Characterising the native range of *C. carassius* has been hampered in the past, largely due to morphological confusion with closely related species (Wheeler 2000; Hickley & Chare 2004). *C. carassius* is presently assumed to be native in southeast England on the basis of two pieces of evidence. Firstly, he identification of *C. carassius* pharyngeal bones found at a Roman archaeological dig site Southwark, London (Lever 1977; Jones 1978), and secondly the similarity of its distribution, in southeast England, to those of other native freshwater fish species, such as silver bream, *Blicca bjoerka* (L.), Ruffe, *Gymnocephalus cernuus* (L.), burbot *Lota lota* (L.) and spined loach, *Corbitis taenia* (L.) (Wheeler 1977, 2000). However, in contrast, Maitland (1972) suggested that *C. carassius* was introduced to south east England along with common carp in the 15^th^ century. More recently, Jeffries et al. (2015) inferred substantial shared ancestry between UK and several Belgian and German populations from microsatellite and genome wide SNP markers supporting the hypothesis of a more recent origin.

Recently, Approximate Bayesian Computation (ABC) methods have been developed (Cornuet *et al.* 2008), that allow such questions to be addressed more explicitly in a population genetic framework, which is suitable for investigating events on a post-Pleistocene timescale. In the present study, we employ ABC to empirically test the status of *C. carassius* in southeast England, using highly polymorphic microsatellite markers. Specifically we test three possible alternative hypotheses for the *C. carassius* colonisation of England; i) all English populations originate from natural colonisation from Continental Europe more than 7800 YBP, ii) all English populations were introduced by humans from Continental Europe sometime in the last 2000 years or iii) some English populations are native and some have been more recently introduced. Our ultimate aim is to increase the knowledge available for the assessment of status and conservation of *C. carassius* in England and Continental Europe.

## Methods

### Samples, DNA extraction and microsatellite amplification

The samples used in this study include 257 *C. carassius*, from 11 English populations, three Belgian populations and one German population (Table 1, Figure 1). These represent a subset of samples from a Europe-wide phylogeographic study, which used the same 13 microsatellite loci as used here, as well as mitochondrial DNA sequences and genome wide SNP data (see Jeffries et al 2015 for Methods). In Jeffries et al 2015, population structure analyses of the Europe-wide dataset showed that these fall into a single genetic cluster, which was distinct from the other genetic clusters found in Europe. The Belgian and German samples used in the present study therefore represent the closest known relatives of English *C. carassius* populations in Europe (Jeffries et al 2015) and are the most likely of our sampled populations to have been the source of their colonisation.

**Table 1.** Location, number and summary statistics of samples used in the present study for microsatellite analyses.

| Code | Location | Country | Drainage | Coordinates |  | N | H<sub>obs</sub> | A<sub>r</sub> |
| --- | --- | --- | --- | --- | --- | --- | --- | --- |
|  |  |  |  | lat | long |  |  |  |
| GBR1 | London | U.K. | U.K | 51.5 | 0.13 | 9 | 0.11 | 1.33 |
| GBR2 | Reading | U.K. | U.K | 51.45 | -0.97 | 4 | 0.03 | NA |
| GBR3 | Norfolk | U.K. | U.K | 52.86 | 1.16 | 7 | 0.16 | 1.48 |
| GBR4 | Norfolk | U.K. | U.K | 52.77 | 0.75 | 27 | 0.12 | 1.26 |
| GBR5 | Norfolk | U.K. | U.K | 52.77 | 0.76 | 14 | 0.13 | 1.30 |
| GBR6 | Norfolk | U.K. | U.K | 52.54 | 0.93 | 20 | 0.22 | 1.55 |
| GBR7 | Norfolk | U.K. | U.K | 52.9 | 1.15 | 24 | 0.15 | 1.44 |
| GBR8 | Hertfordshire | U.K. | U.K | 52.89 | 1.1 | 37 | 0.16 | 1.43 |
| GBR9 | Norfolk | U.K. | U.K | 52.8 | 1.1 | 27 | 0.09 | 1.27 |
| GBR10 | Norfolk | U.K. | U.K | 52.89 | 1.1 | 14 | 0.21 | 1.69 |
| GBR11 | Norfolk | U.K. | U.K | 52.92 | 1.16 | 20 | 0.18 | 1.55 |
| BEL1 | Bokrijk | Belgium | Scheldt River | 50.95 | 5.41 | 13 | 0.15 | 1.42 |
| BEL2 | Meer van Weerde | Belgium | Scheldt River | 50.97 | 4.48 | 12 | 0.19 | 1.48 |
| BEL3 | Meer van Weerde | Belgium | Scheldt River | 50.97 | 4.48 | 8 | 0.16 | 1.47 |
| GER2 | Münster | Germany | Rhine River | 51.89 | 7.56 | 21 | 0.4 | 2.37 |

**Figure 1.**
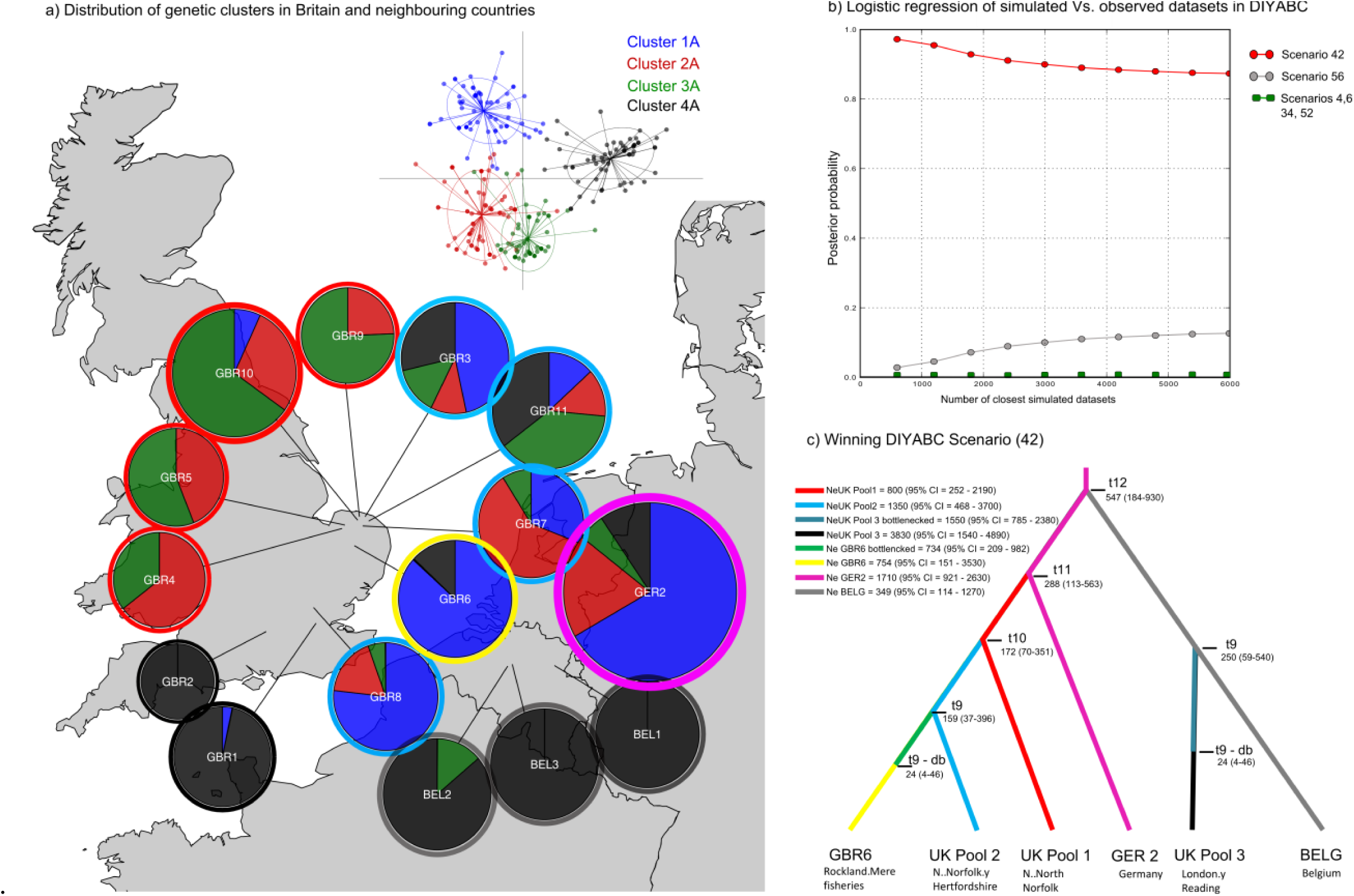
a) DAPC analysis of C. carassius in northwest Europe, showing similar genetic composition of English and Continental populations; b) Posterior probabilities that each of the of the 6 likely DIYABC scenarios explains the distribution of diversity in the northwest European C. carassius, calculated using linear regression between the observed dataset and the closest 6000 simulated datasets; c) Scenario 42 - the winning DIYABC, in which C. carassius were brought to the UK approx. 288 generations ago (t11).

DNA was extracted from tissue samples using either the Puregene DNA isolation kit or the DNeasy DNA purification kit (Qiagen, Hilden, Germany). Samples were then genotyped at 13 microsatellite loci, which were amplified in three multiplex reactions using the Qiagen multiplex PCR mix with manufacturer’s recommended reagent concentrations, including Q solution and 1 µl of template DNA. The annealing temperature was 54°C for all reactions and individual primer pair concentrations within each multiplex reaction were optimised depending on the relative PCR product yield for each locus (see Chapter 3). PCR reactions were run on an Applied Biosciences® Veriti Thermal Cycler and microsatellite fragment lengths were analysed on a Beckman Coulter CEQ 8000 genome analyser using a 400 bp size standard.

### Standard Population statistics

First, allele dropout and null alleles in the data were tested for using Microchecker (Van Oosterhout *et al.* 2004). FSTAT v. 2.9.3.2 (Goudet 2001a) was then used to check for linkage disequilibrium (LD) between loci, deviations from Hardy-Weinberg equilibrium (HWE) within populations and for all population genetic summary statistics. Genetic diversity within populations was estimated using Nei’s estimator of gene diversity (*H*_e_) (Nei 1987) and Allelic richness (*A*_r_), which was standardised to the smallest sample size (n =7) using the rarefaction method (Petit *et al.* 1998). In order to quantify differentiation among populations, pairwise *F*_ST_ values were calculated in FSTAT (Goudet 2001b) using the multilocus (Weir & Cockerham 1984) *F*_ST_ estimator. Sequential Bonferroni correction and permutation tests (2100 permutations) were used to test for significance of *F*_ST_. We also used the Hierfstat package (Goudet 2005) in R (R Core Team 2013), to quantify the genetic variation (*F*_ST_) at 4 hierarchical levels of population isolation, the population-level (separate ponds within countries), the country-level (between Belgium and Germany) the landmass-level (between England and continental Europe) and also at the level of the DIYABC pools used (described below). In the latter case, hierarchical *F*_ST_s were used to validate the population poolings used for the DIYABC as in Pedrischi et al. (2013)

### Testing the native status of C. carassius in England

In order to test our three alternative hypotheses for the colonisation of *C. carassius* in England, an Approximate Bayesian Computation (ABC) approach was taken, implemented in the program DIYABC (Cornuet *et al.* 2014). DIYABC simulates datasets of expected summary statistics (ESS) for user-defined demographic scenarios (‘scenario’ is used herein to describe a specific population tree topology together with the parameter distribution priors that are associated with it). These scenarios were then statistically compared to the actual observed data, allowing us to identify those that are most likely to represent the true history of populations (Cornuet *et al.* 2008).We then estimated the divergence time between populations based on posterior parameter distributions to provide a likely date for the arrival of *C. carassius* in the UK.

In order to reduce the number of scenarios to be tested from the huge number possible, we grouped populations in DIYABC analyses into pools of populations with shared history, a method also employed by Pedreshi et al. (2013). To inform these poolings it was first necessary to perform a fine scale population structure analysis of the 15 populations used. This was done using Discriminant Analyses of Principal Components (DAPC), implemented in the Adegenet R package (Jombart *et al.* 2010). Bayesian Information Criteria (BIC) scores were used to choose the appropriate number of genetic clusters in the dataset. Spline interpolation (Hazewinkel 1994) was then used to identify the appropriate number of principal components for use in the subsequent discriminant analysis.

Based on the results of the DAPC analysis, populations were grouped into six pools. Those of similar genetic composition (and therefore very likely to have a shared history) were pooled together (see results section). However, if populations from either side of the English Channel shared similar genetic composition, they were separated across pools, to allow for hypothesis testing.

In total, 56 scenarios were tested: six, 39 and 11 representing hypothesis i), ii) and iii) respectively (Supplementary Figure 1). The number of scenarios for each hypothesis reflects the number and plausibility of the possible population histories for the different hypotheses given the results of the populations structure analysis. The discriminating factors between scenarios representing different hypotheses were tree topology and, most importantly, the parameter priors for the divergence times between populations (Supplementary table 1). These divergence time priors were set in order to represent the possible time windows of *C. carassius* introduction under our three hypotheses. To test hypothesis i) – the natural colonisation of *C. carassius* more than 7800 YBP - the time prior for the oldest split between English and Continental European populations was set to 4000-10000 generations (equivalent to 8000 – 20000 YBP, assuming an average generation time of two years (Tarkan et al. 2010), Supplementary Figure 1: scenarios 1-6). To test hypothesis ii) – that English *C. carassius* were introduced after the 15^th^ century - the same prior was set to 10-1000 generations (2 – 2000 YBP, scenarios 25 - 44), which very conservatively encompasses all dates of possible live fish translocations to the UK by humans. Finally, to test hypothesis iii) - that some populations were native and some introduced we used multiple combinations of both native and introduced prior dates (as used in hypothesis i and ii) scenarios respectively) for different population splitting events (scenarios 45 – 56). In the interests of completeness, we also tested an intermediate time window of 10 – 2500 generations (20 – 5000 YBP, scenarios 7-24). Analyses were performed in a sequential manner, whereby a million datasets per scenario were first simulated in DIYABC. Then, for the computationally intensive part of the analysis, simulated datasets were grouped according to the hypothesis they represented (i.e. (i), (ii) or (iii)) and these groups were separately compared to the observed data using both approaches offered in DIYABC, logistic regression and “direct estimate”. The latter of which is a count of the number of times that a given scenario simulates one of the closest datasets to the real data set (Cornuet *et al.* 2008). The resulting posterior probabilities were used to identify the top two most likely scenarios for each hypothesis (six in total). These were then used in a final test, again using logistic regression and direct estimate, to identify the single most likely scenario of the final 6. Model checking analyses, which measures the discrepancy between the model parameter posterior combination and the actual data (Cornuet *et al.* 2010), were then carried out to test the robustness of scenario choice. Finally, posterior parameter distributions for effective population size, divergence times and bottleneck parameters were estimated on the basis of the most likely scenario.

## Results

### Microsatellite data analyses

Microchecker showed no consistent signs of null alleles or allele dropout in populations of pure *C. carassius* and no LD was found between loci pairs. Tests of Hardy-Wienberg proportions did not identify any populations that significantly deviated from HWE.

### Population Structure in England, Belgium and Germany

Population structure was weakest (0.0) between the two Belgian populations, strongest (0.736) between GBR2 and GBR4 (Supplementary table 2) and followed a weak IBD pattern, being significantly associated with geographic distance (adjusted *R*^2^ = 0.248, *P* < 0.001, Supplementary Figure 2). Hierarchical assessment of population structure showed that variation between individuals was significantly explained by population assignment and country (F_pop_ *=* 0.36, *P =*0.001; F_country_ = 0.154, *P =*0.001). However the landmass (continental Europe or Britain) had no significant effect on variation between individuals (F_landmass_= -0.04, *P =*0.482). Importantly, the pools used in DIYABC analysis explained a large of the genetic between individuals in total (F_pools_ = 0.244, *P =*0.001) and within the poolings the remaining variation between individuals was considerably lower than at the landmass level or the country level, though still highly significant (F_Ind/pools_ = 0.142, *P* = 0.001), confirming that these population groupings were appropriate groupings for the populations in DIYABC analyses.

Observed heterozygosity (averaged across all loci within a population) ranged from 0.03 (GBR2) to 0.4 (GER2). A_r_ ranged from 1.26 (GBR4) to 2.37 (GER2), and correlated with H_o_ (adjusted *R*^2^ = 0.543, *P* = 0.001).

In the DAPC analysis of population structure, ten genetic clusters were indicated by BIC scores (Supplementary Figure 3c). The resulting population-cluster identities were complex (Supplementary Figure 3b), with most populations containing many closely related clusters (Supplementary Figure 3a) making it difficult to identify sets of closely related population for pooling. Therefore in order to reliably inform our DIYABC poolings, we incrementally dropped the number of clusters to four which seem to reflect the large scale patterns of genetic differentiation better. Seven principal components and two linear discriminants were retained in this final, four-cluster DAPC analysis (Figure 1a). The resulting inferred population structure showed that many of the English populations showed higher similarities to Continental populations than to neighbouring English populations. For example, GBR1 and GBR2 were extremely similar to Belgian populations, and GBR3, 6, 7, 8 and 11 were more similar to populations in northern Germany (Figure 1a). However, GBR4, 5, 9, 10, all in north Norfolk (eastern England), showed some distinctiveness from continental populations.

### Testing the native status of C. carassius in England

For the DIYABC analyses, populations were grouped into six pools on the basis of the above DAPC results (pools are denoted by coloured rings around pie charts in Figure 1a). Within-hypothesis logistic regressions of simulated vs. observed data, performed in DIYABC, showed that the two most likely scenarios for each hypothesis were scenarios 4 and 6 for hypothesis i); 42 and 34 for hypothesis ii) and 52 and 56 for hypothesis iii). These final six scenarios were then tested against each other, again using logistic regression to find the single most likely scenario of all 56 tested. Scenario 42, representing hypothesis ii), produced data sets that were, by far, the closest to the real data, with a posterior probability of 0.91 (Figure 1b).

Scenario 42 (Figure 1c) had prior constraints on the split between English and Continental populations (t11) of 10 – 1000 generations and thus supports a human introduction of *C. carassius* into southeast England <2000 YBP. Under this scenario, the oldest demographic event was the split between German and Belgian populations approximately 547 generations ago (1094 YBP). However, the most important demographic event for the purposes of testing our hypotheses is the split between English populations (UK pools 1, 2 and RM) and continental populations (pools GER2 and BELG), at time “t11” in Scenario 42 (Figure 1c). Furthermore, this scenario suggests that the ancestral source population of the initial English introduction was more closely related to the German than the Belgian populations sampled here. The date of this English/Continental population split is estimated at 288 (95% CI = 113-563, Supplementary Table 3) generations ago, which corresponds to 576 (95% CI = 226 - 1126) YBP, approximately 7400 years after the loss of the Doggerland land bridge. DIYABC also outputs posterior estimates of population split times scaled by mutation rate and effective population size. The estimated time for the English/Continental population split, scaled by mutation rate estimated by the model was t11(u+_SNI_) = 9.83 × 10^−2^ (where u+SNI is the median estimate of the microsatellite mutation rate using the generalised stepwise mutation model, (1.11×10^−4^mutations/locus/generation) and SNI is the single nucleotide insertion rate (6.18×10^−8^/mutations/locus/generation) Supplementary table 3). The median estimate of this mutation rate (u = 1.11×10^−4^ /locus/generation), although slow, is still within the realms of that observed in the closely related *C. carpio* (mean = 5.56×10^−4^ mutations/locus/generation, 95% CI = 1.52×10^−4^ - 1.63×10^−3^, (Yue *et al.* 2007)) and indeed in humans (Ellegren 2004).

To validate this result we first tested the “goodness-of-fit” of Scenario 42 using statistical model checking as implemented in DIYABC, which showed that the observed data fell well within the predictive posterior parameter distribution of the simulated data (Supplementary Figure 4). Secondly, we calculated the oldest possible date of the English/Continental population split using its upper 95% confidence value under Scenario 42 (563 generations), and assumed the unrealistic, but sometimes possible generation time of 5 years (Tarkan *et al.* 2010). Despite these extremely conservative values, the split between English and Continental populations was still estimated at 2815 YBP, approximately 5000 years after the flooding of Doggerland. Finally, we inferred t11 (the English/Continental population split) of scenario 42 using the scaled parameter estimate, t11(u+SNI). This gave an estimate of 885 generations, or 1770 years (with a two year generation time), which, although older than the un-scaled estimate, is still over 6000 years later than the possible natural colonisation window. In fact, in order for the scaled estimate to fit the hypothesis of natural colonisation (more than 8000 years ago), assuming a two year generation time, the mutation rate would have to be approximately 1.0×10^−5^mutations/locus/generation, at least one order of magnitude lower than reported for microsatellite loci (reference).

Further population splits have occurred more recently from this initial introduction, and there is also support for a second independent introduction of *C. carassius* into the UK (t9) approximately 250 (95% CI = 59-540) generations or 500 (95% CI = 118-1080) years ago (UK pool 3), from a source population closely related to the Belgian populations sampled here.

## Discussion

The primary aim of the present study was to test the contentious assumption that *C. carassius* arrived in southeast England naturally. Owing to its hydrogeological history during the last glaciation, the UK presents a rare opportunity to test such a question amongst its inhabitants. Our analyses suggest that *C. carassius* was anthropogenically introduced into England and on this basis we therefore discuss the potential implications for *C. carassius* conservation.

### Non-native origins of C. carassius in England

Analyses of the population structure within southeast England and closely neighbouring countries revealed that many English populations are more similar genetically to continental populations than to their English counterparts, implying multiple independent colonisation events or introductions into England. DIYABC analyses supported this, suggesting that populations GBR1 and GBR2 split from Belgian populations more recently than they did from other English populations (Figure 1c). Indeed these populations are known to be managed and therefore have likely been stocked in the recent past; GBR1 being a conservation pond, and GBR2 a fish farm. Therefore, our results indicate that these fish came from recently imported stocks closely related to the sampled Belgian populations.

In contrast to GBR1 and GBR2, DIYABC analyses suggest that all north Norfolk and Hertfordshire populations share a most recent common ancestor with the sampled German population; indicative of a separate introduction. The central question of this analysis however, was; how long ago was the first colonisation or introduction of *C. carassius* into England? DIYABC analyses predicted that the oldest possible date for the arrival of *C. carassius* in England was approximately 1126 YBP but most likely 576 YBP; over 7000 years after the loss of the Doggerland land bridge, and that there were in fact two independent introductions around this time.

As this result could have important implications for the conservation of *C. carassius* in the UK (see below), we performed rigorous results checking. Tests for the goodness-of-fit of the winning scenario (42) confirmed that this was the most likely out of all scenarios tested, and even when using the 95% confidence interval limits of the posterior time parameter distribution or using the unrealistically long generation time of 5 years (to convert DIYABC results from generations to years), it still was not possible to achieve estimates of the split between English and continental populations older than 2815 YBP. Only with a mutation rate an order of magnitude slower than that estimated here (and elsewhere, e.g in *C. carpio* (Yue *et al. 2007), mice (Dallas 1992), sheep (Crawford & Cuthbertson 1996) and humans (Ellegren 2004)) would the time for this split support a natural introduction of C. carassius* into England.

Although our sampling is not exhaustive, it comprehensively covers the areas of England previously thought to contain native *C. carassius* populations, in particular Norfolk, which is thought to have been a stronghold for *C. carassius* in the past (Patterson 1905; Ellis 1965; Sayer *et al.* 2011). It is therefore unlikely that there are unsampled populations of *C. carassius* in England that show further divergence from those of continental Europe. Furthermore, broad scale phylogeographic results in Jeffries et al (2015) show that Belgian and German populations are the closest relatives of English *C. carassius* in Europe. In fact, adding currently unsampled populations from continental Europe could only result in a lower estimate of divergence between English and continental European samples. We are, therefore, confident that our estimate represents the earliest possible timeframe for the first *C. carassius* introductions into England. It should also be noted that the estimate for this split does not directly predict when populations were introduced to England, only when they were separated from the sampled continental European populations, which must have been at the same time as, or prior to, their introduction. Thus, it is entirely possible that the arrival time of *C. carassius* in the UK was even more recent than the DIYABC estimate of population divergence time.

However, we cannot rule out the possibility that *C. carassius* colonised naturally, but either then went extinct, or were extirpated by the current English *C. carassius* strains when they were introduced. If these scenarios were true, only dated fossil evidence, and perhaps ancient molecular studies would allow for a definitive answer.

The results of this study therefore strongly point to the anthropogenic introduction of English *C. carassius* and, in fact, fall perfectly in line with the first known record of *C. carpio* introductions into England by monks for food in the 15^th^ Century (Lever 1977). However, we can only speculate as to the motivations behind these introductions. To our knowledge, *C. carassius* are not mentioned in the literature until 1766 (Pennant 1766), however it is possible that *C. carassius* was intentionally introduced as a source of food, as with *C. carpio*. Indeed there are mentions of *C. carassius* used as food in 1778 in Norfolk (Woodforde *et al.* 2008), and although *C. carassius* does not grow to the size of other carp species, its ability to survive in small, isolated and often anoxic ponds may have made it an attractive species for use in medieval aquaculture. It is possible, however, that the introduction of *C. carassius* in England was unintentional. For example, it can be very difficult to tell *C. carassius* and *C. carpio* apart, especially if they are found in sympatry and if hybrids are present (Wheeler 2000), as is often the case (Hänfling *et al.* 2005; Sayer *et al.* 2011). Irrespective of the initial motivations however, intentional movements of *C. carassius* have since been common, predominantly for angling purposes (Sayer *et al.* 2011).

### Conclusions and implications for the conservation of C. carassius

A fundamental consideration in the conservation of a species is its native range, and, contrary to current belief, the results of this study support the human-mediated introduction of *C. carassius* into England. But what does this mean for the conservation of *C. carassius* in England, a country which has one of the few active projects in place for its conservation (Copp & Sayer 2010)? In light of these results, should England cease efforts conservation of *C. carassius*? There has been a call recently, for a change in the conservation paradigm, moving away from the unfounded assumption that all non-native species have detrimental impacts on native ecosystems (Davis *et al.* 2011). Instead the authors advocate embracing the idea of constantly changing communities, and moving towards impact-driven conservation, whereby only those species that have been empirically shown to be invasive and detrimental to native ecosystems and economies are actively managed. Indeed only a small proportion of freshwater fish introductions have been shown to have detrimental impacts on the native ecosystem, whereas many provide significant ecological and economical benefits (Gozlan 2008; Schlaepfer *et al.* 2011), and sometimes replace ecosystem services lost in extinct species (Schlaepfer *et al.* 2011). Currently, *C. carassius* could not be labelled as invasive in England, as they are not expanding, in fact, they are declining in numbers in England (Sayer *et al.* 2011). To date, there has been no attempt to assess the impact of *C. carassius* on ecosystems due to the assumption that they were native, however, available studies show that *C. carassius* are widely associated with species-rich, macrophyte-dominated ponds (Sayer *et al.* 2011), which are extremely important ecosystems for conservation (Oertli *et al.* 2002). There is no evidence that *C. carassius* negatively impact these habitats, unlike *C. carpio (Miller & Crowl 2006), and despite concerns that C. carassius* may impact the threatened great crested newt (*Triturus cristatus,* Laurenti 1768), this does not seem to be the case in UK ponds, with *C. carassius* often co-existing with recruiting *T. cristatus* populations (Chan 2010).

A further important consideration in the case of *C. carassius* is its threatened status in much of its native European range. Copp *et al.* (2005) *pose the question; should we treat all introduced species in the same way, even if one such species is endangered in its native range? Indeed, if the goal of conservation science is to protect and enhance biodiversity, it would seem counterproductive to abandon the conservation of C. carassius* populations in one region when they are threatened in another. Our Europe-wide population structure results show that English populations, along with those in Belgium and Germany, comprise a distinct part of the overall diversity of *C. carassius* in Europe. And this is made all the more important by the expansion of *C. gibelio* through Europe, especially into the Baltic Sea basin from the south (Wouters *et al.* 2012; Deinhardt 2013); Lauri Urho. Pers. comms). Although the invasive *C. auratus* is present and poses a threat to *C. carassius* in England (as it does in continental Europe), *C. gibelio* is not yet present and therefore England may represent an important refuge from this threat.

A final consideration for the continued conservation of *C. carassius* is their status as an English heritage species. *C. carassius* is affectionately regarded by the zoological and angling communities of England and as such, has regularly featured in the writings of both groups over the past three centuries (see the many examples in (Rolfe 2010), pp. 50-64). Therefore, although our results indicate that *C. carassius* can probably not be regarded as a native species in the true sense, the species been has been an important part of the cultural landscape in England for around 500 years.

As outlined above, despite the evidence that *C. carassius* is non-native in England, strong arguments can be made for its continued conservation in that important part of its range. However, our results bring to light much broader and timely questions in invasion and conservation biology; how many assumptions about the native status of other freshwater species in the UK would stand up to the same tests as performed here, if the data were available to perform it? And what do we do about it if they don’t?

## Acknowledgements

The authors thank the FSBI (www.fsbi.org.uk) and Cefas (Lowestoft, UK) for funding this research. We also thank all land owners and the following contributors of fish tissue; Keith Wesley, Ian Patmore and Dave Emson (England).

## Supplementary materials

**Supplementary table 1.**
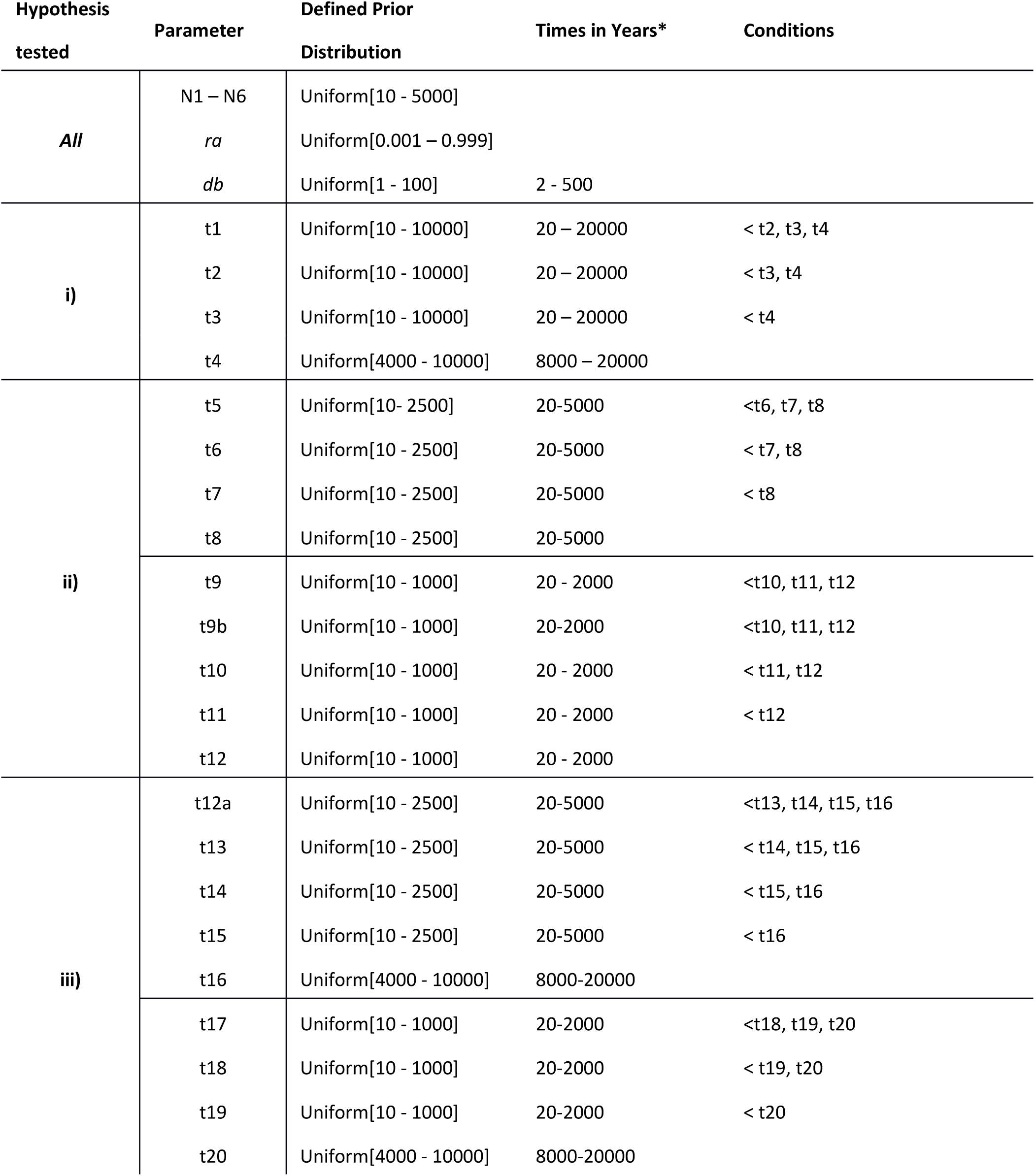
Prior parameters for all scenarios used in DIYABC analyses.

**Supplementary table 2.** Pairwise FST values for 15 C. carassius populations in northwest Europe.

|  | GBR1 | GBR2 | GBR4 | BEL1 | BEL2 | BEL3 | GER2 | GBR7 | GBR3 | GBR8 | GBR9 | GBR11 | GBR5 | GBR6 | HUN3 |
| --- | --- | --- | --- | --- | --- | --- | --- | --- | --- | --- | --- | --- | --- | --- | --- |
| GBR1 | 0 | 0.3148 | **0.6323** | **0.3711** | **0.207** | 0.366 | **0.2842** | **0.5017** | 0.3676 | **0.4369** | **0.6114** | **0.2975** | **0.5956** | **0.3518** | **0.4373** |
| GBR2 |  | 0 | **0.7369** | 0.353 | **0.2644** | 0.3795 | **0.3334** | **0.6109** | 0.535 | **0.5616** | **0.7164** | **0.3809** | **0.7** | 0.4851 | 0.5358 |
| GBR4 |  |  | 0 | 0.6616 | **0.494** | 0.5857 | **0.3076** | **0.1943** | 0.4229 | **0.3497** | **0.2982** | **0.2058** | **0.2351** | **0.4976** | **0.1883** |
| BEL1 |  |  |  | 0 | 0.0755 | 0.0204 | **0.269** | **0.5601** | **0.3835** | **0.4882** | **0.5886** | **0.2985** | **0.5648** | **0.3322** | **0.4858** |
|  |  |  |  |  |  | - |  |  |  |  |  |  |  |  |  |
| BEL2 |  |  |  |  | 0 | 0.0149 | **0.1878** | **0.4225** | **0.2504** | **0.3869** | **0.4159** | **0.157** | **0.3934** | **0.2715** | **0.3002** |
| BEL3 |  |  |  |  |  | 0 | **0.1909** | **0.4879** | **0.3133** | **0.4379** | **0.5058** | **0.2165** | **0.4763** | **0.299** | **0.3754** |
| GER2 |  |  |  |  |  |  | 0 | **0.1669** | 0.0682 | **0.1352** | **0.3144** | **0.134** | **0.2451** | **0.1502** | **0.1845** |
| GBR7 |  |  |  |  |  |  |  | 0 | 0.1526 | **0.0717** | **0.3643** | **0.164** | **0.2876** | **0.3282** | **0.161** |
| GBR3 |  |  |  |  |  |  |  |  | 0 | 0.0208 | **0.422** | 0.0896 | **0.3539** | 0.0795 | 0.2133 |
| GBR8 |  |  |  |  |  |  |  |  |  | 0 | **0.4205** | **0.1837** | **0.3569** | **0.2124** | **0.2581** |
| GBR9 |  |  |  |  |  |  |  |  |  |  | 0 | **0.2054** | 0.0287 | **0.4394** | **0.2007** |
| GBR11 |  |  |  |  |  |  |  |  |  |  |  | 0 | **0.1633** | **0.2274** | **0.1071** |
| GBR5 |  |  |  |  |  |  |  |  |  |  |  |  | 0 | **0.3894** | **0.1972** |
| GBR6 |  |  |  |  |  |  |  |  |  |  |  |  |  | 0 | **0.318** |
| HUN3 |  |  |  |  |  |  |  |  |  |  |  |  |  |  | 0 |

P-values obtained after:2100 permutations
Indicative adjusted nominal level (5%) for multiple comparisons is: 0.000476

**Supplementary table 3.**
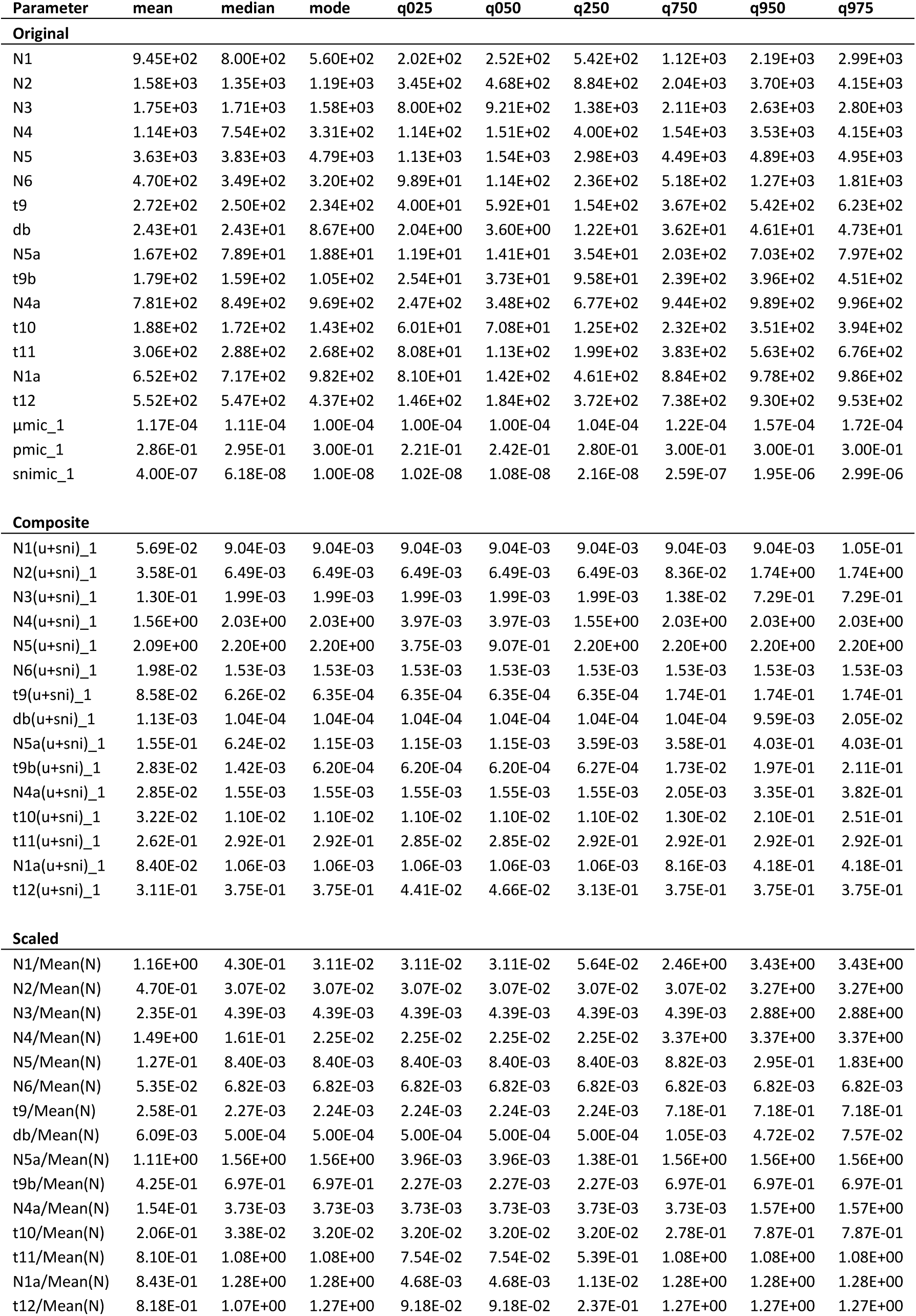
All posterior parameter distributions for all scenario 42 - identified as the most likely scenario for the colonisation of C. carassius into England by DIYABC analyses.

**Supplementary Figure 1.**
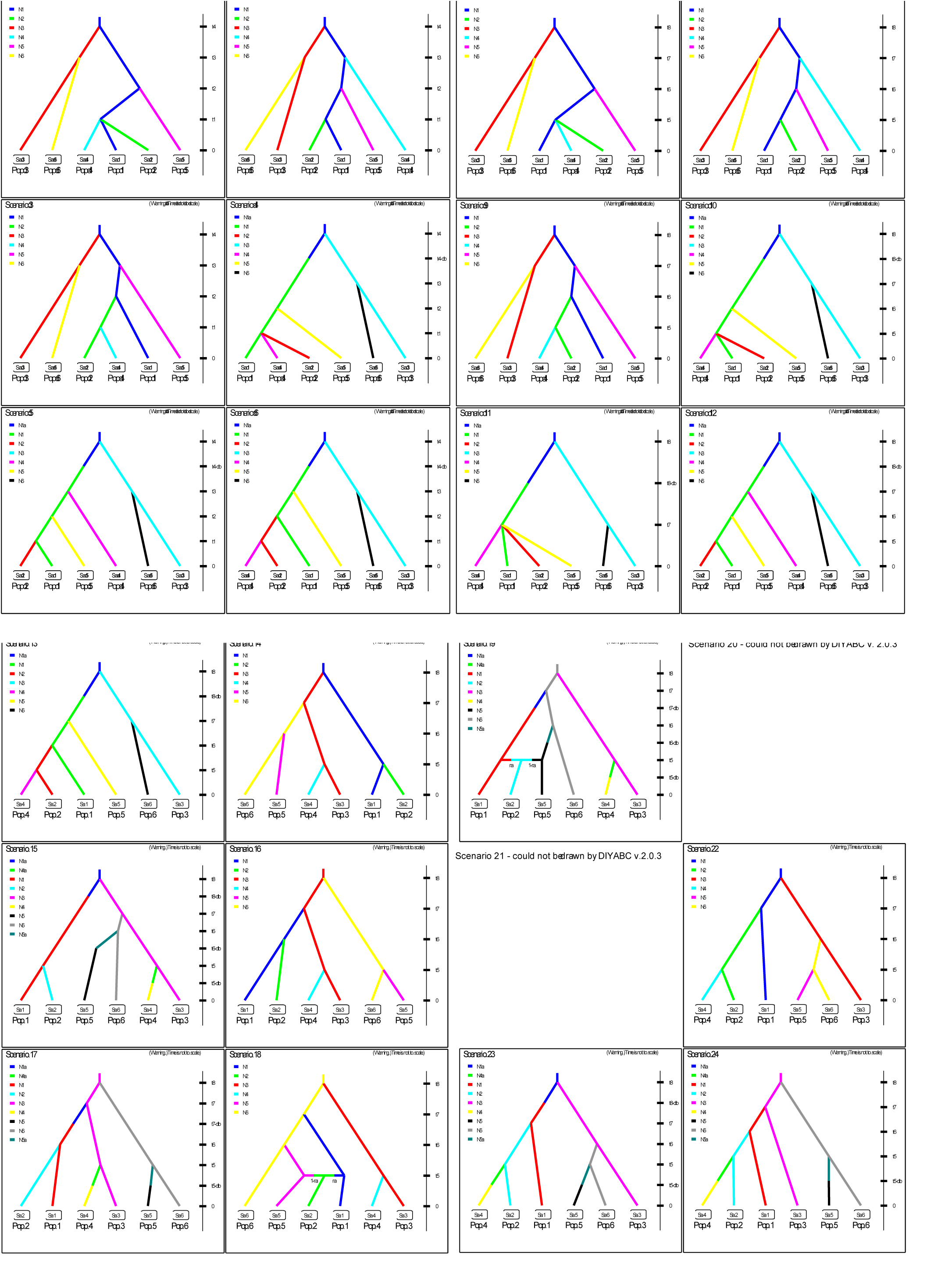
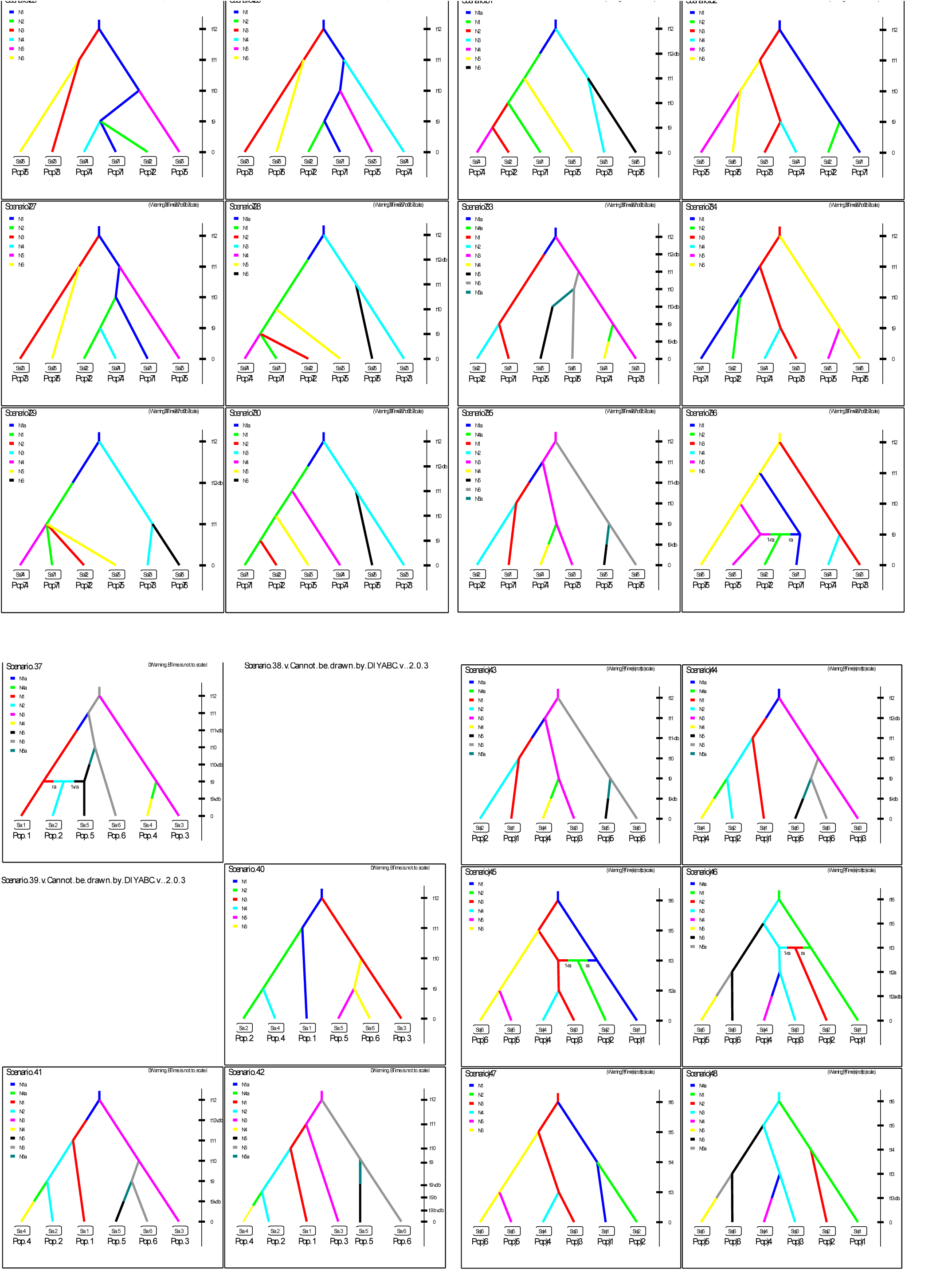
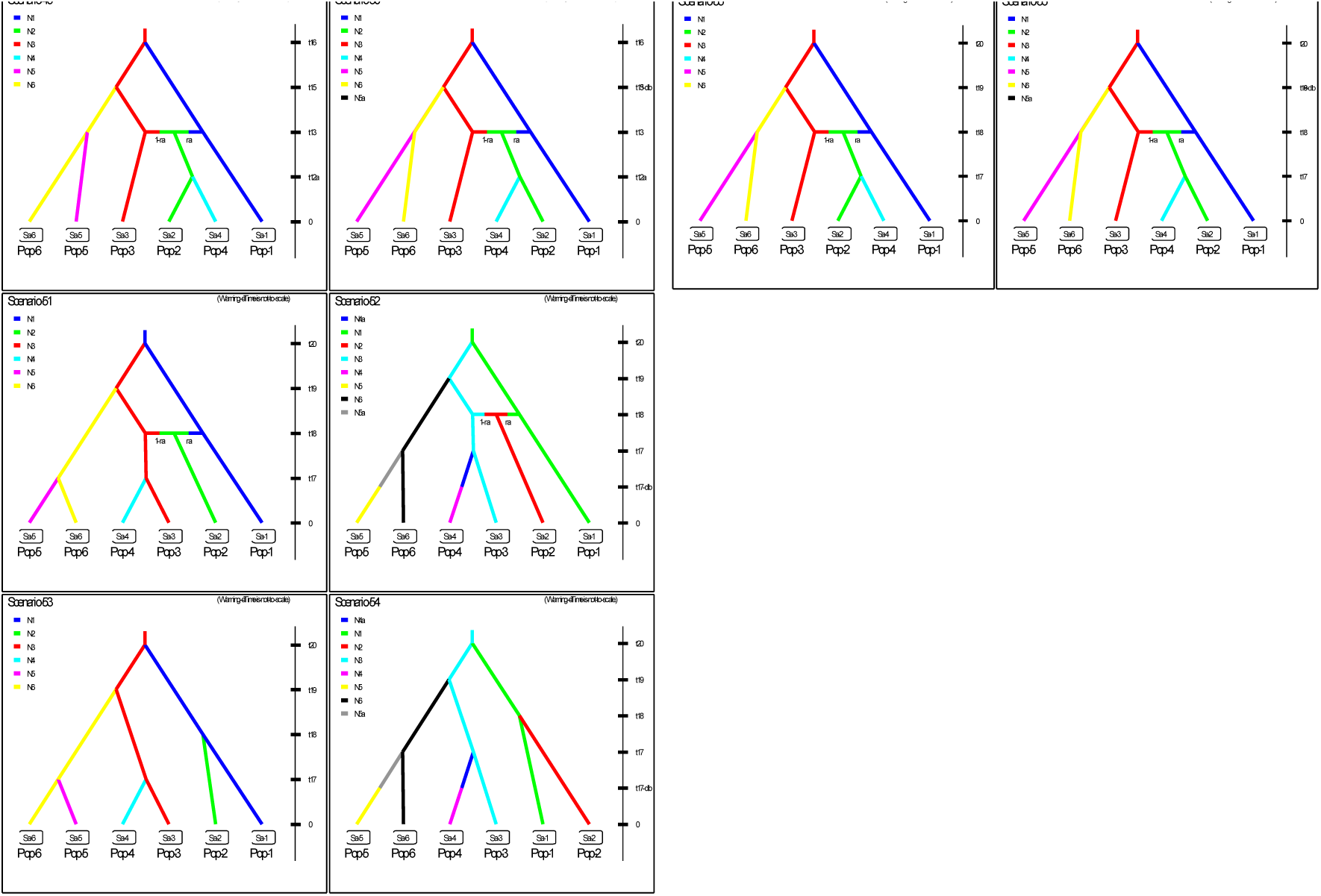
All scenarios tested in DIYABC analysis. Pop1 = UK Pool 1, Pop2 = UK Pool 2, Pop3 = GER, Pop4 = UK pool 4, Pop5 = UK Pool 3, Pop6 = BELG. For the user-defined prior parameter distributions see S.Table 2.

**Supplementary Figure 2.**
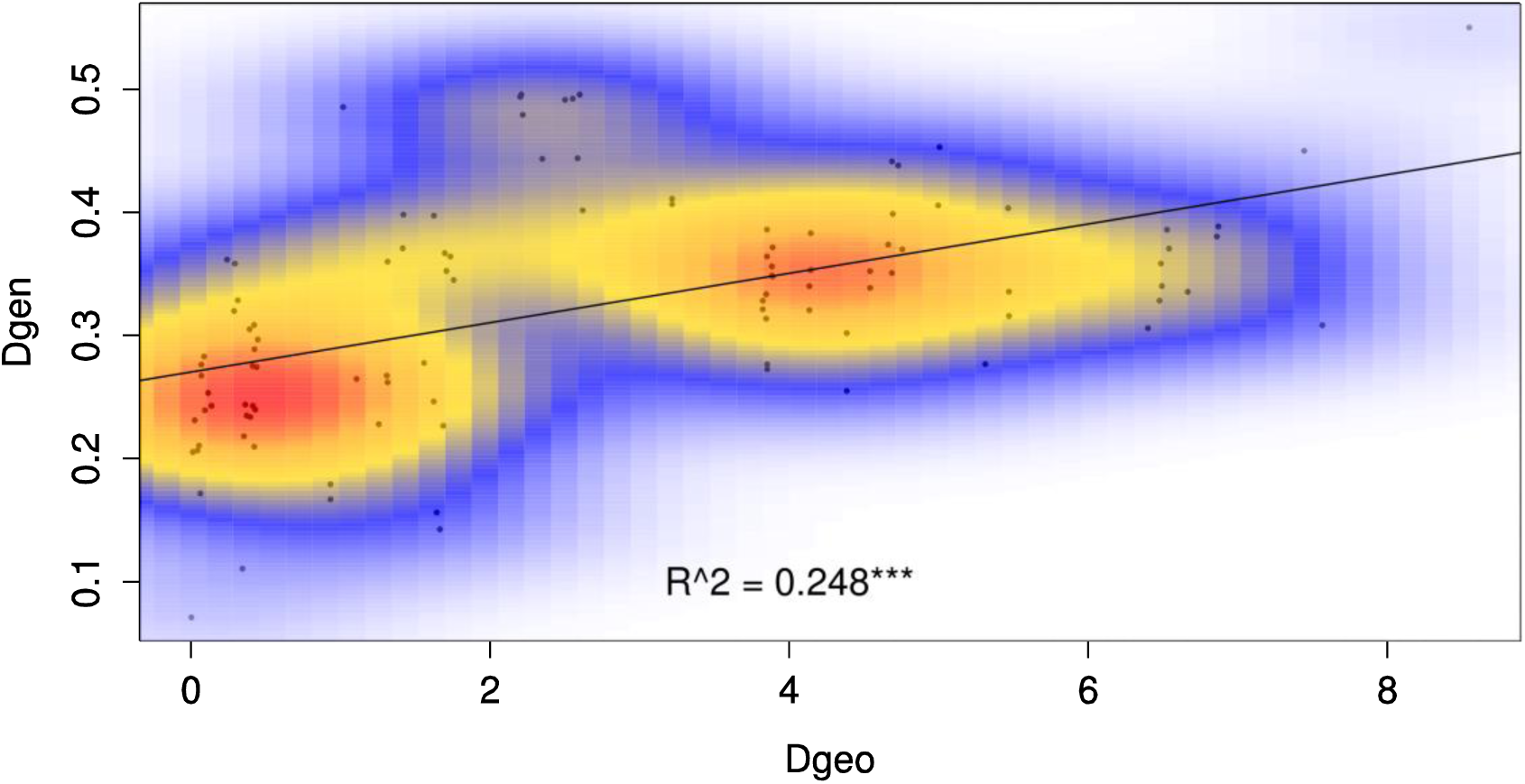
Isolation by distance in the 15 populations sampled.

**Supplementary Figure 3.**
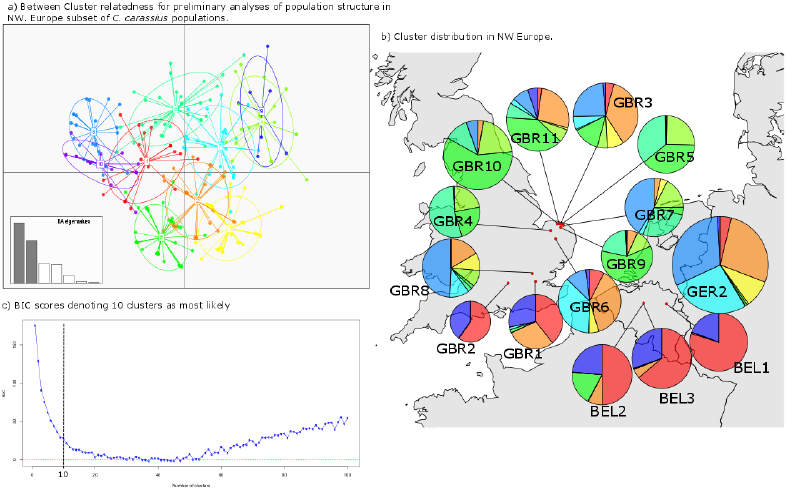
DAPC analysis of English, German and Belgian C, carassius populations. a) Shows relatedness between inferred clusters, b) shows geographic distribution of those clusters within populations and c) gives the BIC scores denoting 10 clusters as the most likely (the number of clusters after which no significant change in BIC score is observed).

**Supplementary Figure 4.**
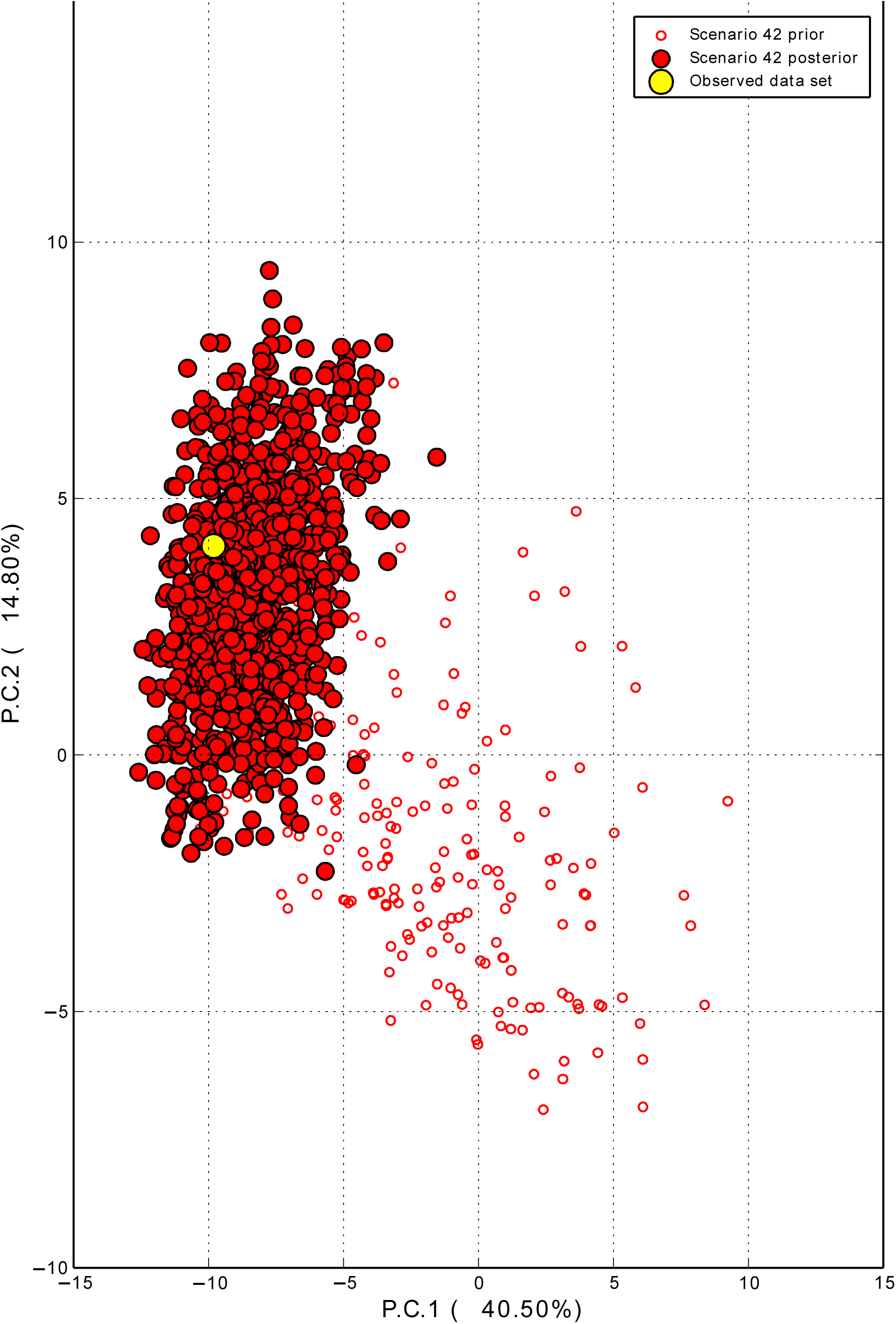
The results of Model Checking of the most likely scenario identified in DIYABC. Note that Observed dataset lies well within the cloud of the predictive posterior parameter distribution.

